# Nonsense-mediated decay is highly stable across individuals and tissues

**DOI:** 10.1101/2021.02.03.429654

**Authors:** Nicole A. Teran, Daniel Nachun, Tiffany Eulalio, Nicole M. Ferraro, Craig Smail, Manuel A. Rivas, Stephen B. Montgomery

**Affiliations:** Department of Pathology, School of Medicine, Stanford University, Stanford, CA, USA; Department of Genetics, School of Medicine, Stanford University, Stanford, CA, USA; Biomedical Informatics Program, Stanford University, Stanford, CA, USA; Genomic Medicine Center, Children’s Mercy Research Institute and Children’s Mercy Kansas City, Kansas City, MO 64108, USA; Department of Biomedical Data Science, Stanford University, Stanford, CA, USA

## Abstract

Precise interpretation of the effects of protein-truncating variants (PTVs) is important for accurate determination of variant impact. Current methods for assessing the ability of PTVs to induce nonsense-mediated decay (NMD) focus primarily on the position of the variant in the transcript. We used RNA-sequencing of the Genotype Tissue Expression v8 cohort to compute the efficiency of NMD using allelic imbalance for 2,320 rare (genome aggregation database minor allele frequency <=1%) PTVs across 809 individuals in 49 tissues. We created an interpretable predictive model using penalized logistic regression in order to evaluate the comprehensive influence of variant annotation, tissue, and inter-individual variation on NMD. We found that variant position, allele frequency, including ultra-rare and singleton variants, and conservation were predictive of allelic imbalance. Furthermore, we found that NMD effects were highly concordant across tissues and individuals. Due to this high consistency, we demonstrate *in silico* that utilizing peripheral tissues or cell lines provides accurate prediction of NMD for PTVs.

## Introduction

RNA expression is not only regulated by transcription but also by degradation (Pai et al. 2012); RNA transcripts with protein-truncating variants (PTVs) are often targeted for degradation by the nonsense-mediated decay (NMD) pathway (Kurosaki, Popp, and Maquat 2019). The accurate identification of PTV-harboring transcripts that are successfully cleared by NMD can have a large effect on disease outcome. Some nonsense mutations lead to dominant-negative effects where the truncated allele can impede the function of the full length allele (Khajavi, Inoue, and Lupski 2006). Mendelian disease diagnostics can benefit from the identification of the PTVs that escape NMD and may therefore lead to truncated peptides and corresponding gain-of-function effects (Coban-Akdemir et al. 2018). To be able to improve identification of PTVs that undergo or escape NMD, existing tools have integrated variant-level annotations which provide a prediction of the NMD efficiency, or ability for a PTV containing transcript to be targeted and degraded by the NMD machinery as measured by the relative amount of a PTV containing transcript as compared to the wild-type (Nagy and Maquat 1998; Lindeboom, Supek, and Lehner 2016; Rivas et al. 2015).

Position explains most variation in NMD efficiency, summarized by the 50 nucleotide (50nt) rule: if the variant occurs farther upstream than 55 to 50 nucleotides before the last exon junction, it will be targeted for degradation. Additional analysis in cancer has indicated that falling near the start of a gene or in a long exon (>407 base pairs) impedes degradation and a simple decision tree, called NMDetective-B, which utilizes these rules can explain 68% of the variation in NMD efficiency (Lindeboom et al. 2019).

These existing approaches have benefited from measuring NMD effects through allele-specific measurement of RNA-sequencing (RNA-seq) read counts overlying PTV variants. However, there is evidence that the ratio of the RNA read counts from the aberrant allele to that of the wild type allele can vary between tissues, which would not be expected if variant position was the only determining factor (Rivas et al. 2015), (Zetoune et al. 2008).

We utilized the Genotype Tissue Expression (GTEx) dataset to assess the impact of tissue type on NMD efficiency (GTEx Consortium 2020). We measured the functional impact of 2,320 rare (genome aggregation database [gnomAD] minor allele frequency [MAF] <=1%) PTVs from 809 individuals across 49 different tissues. We observed that, in addition to position, allele frequency, including rare, ultra-rare (MAF < 0.001%) and singleton alleles predict NMD efficiency. However, tissue is not predictive of NMD efficiency and PTVs showed more consistent allelic imbalance across tissues than any other type of coding transcript variant. Using this information, we demonstrate that accurate identification of PTVs that either undergo or escape NMD can be further achieved in peripheral tissues or cell lines.

## Results

### Identifying NMD-targeted variants in GTEx

In order to evaluate NMD rules in and between normal human tissues, we annotated the proportion of expressed reference reads for rare PTV sites across the GTEx dataset. We used the genomes of 809 individuals of European descent to identify 2,320 different PTVs with an allele frequency from the gnomAD database less than or equal to one percent. Rare variants were selected in order to prevent inclusion of common variants that appear as false positive PTVs due to selection and adaptation favoring the truncated transcript. The proportion of expressed reference reads was calculated by dividing the number of RNA-seq reads that map to a variant site containing the reference allele by the total number of RNA-seq reads overlapping the variant site in a single sample (i.e. one tissue in one person).

We analyzed RNA sequencing data from 49 distinct tissues where each individual had a median number of 17 tissues and a median of five expressed PTVs that were testable in at least one GTEx tissue. We calculated the proportion of reference reads for each variant in each tissue for a total of 40,402 variant-tissue-subject observations from the 13,849 tissue-subject samples. From these 40,402 observations, 55% (22,301) were predicted to be targeted by the NMD machinery according to rules in Lindeboom *et al*. of not being near the start of a gene, in a long exon, or after 55-50 nucleotides before the last exon junction. The remaining 45% (18,629) of PTVs were predicted to escape NMD. These rules, on a whole, provided good separation of variants that showed allelic imbalance: 52% of observations of variants that were predicted to be targeted by NMD showed allelic imbalance (reference read proportion ≥ 65%) compared to only 20% of those predicted to escape (Figure 1A).

**Figure 1.**
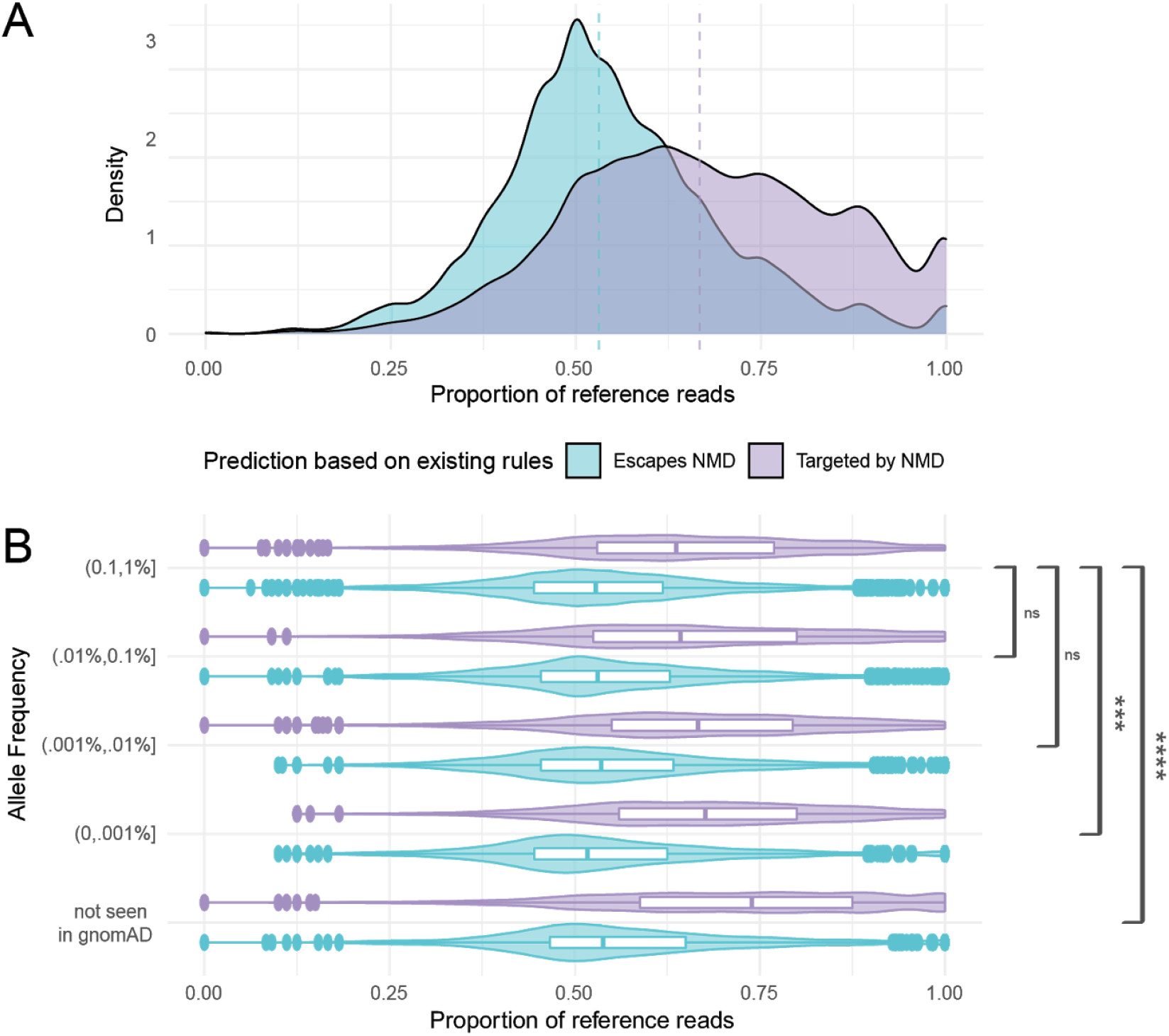
Centrally located and rare truncating variants show stronger allelic imbalance. **A**. Distribution of the proportion of reference reads for rare (genome aggregation database [gnomAD] minor allele frequency [MAF] ≤ 1%) protein truncating variants for those predicted by the positional rules defined in Lindeboom *et al*. to escape NMD (light blue) or trigger NMD (light purple). **B**. Distribution of rare stop variants for variants predicted to escape NMD (light blue) or trigger NMD (light purple) by gnomAD allele frequency. Boxplots show mean and interquartile range. Brackets show the significance of the difference in differences test between each prediction type across decreasing allele frequencies P < 0.00001: ****; P < 0.001: ***; not significant: ns.

### Ultra-rare protein truncating variants have increased allelic imbalance

Previous studies have reported that rarer PTVs are more likely to trigger NMD (Lindeboom et al. 2019; Kukurba et al. 2014; Rivas et al. 2015), including in an earlier version of GTEx (Rivas et al. 2015). This initial exploration of NMD in GTEx analyzed 4,584 PTVs across the allele frequency spectrum acquired from 173 individuals. Given our increased sample size, more extensive whole genome data in GTEx, and the availability of precise allele frequency information from gnomAD (Karczewski et al. 2020), we set out to evaluate this effect with more granularity in the rare allele frequency spectrum. Here, we evaluated the allelic imbalance for PTVs predicted to be NMD targets versus those predicted to be NMD escapees stratified by allele frequency. For rare variants, we saw significant separation between the predicted NMD escapees and the predicted NMD targets; strikingly, this separation was significantly more pronounced (by a difference in differences test) at ultra-rare (MAF < 0.001%) allele frequencies (Figure 1B).

By combining gnomAD allele frequency information with the whole genome sequencing samples from GTEx, we were able to further investigate the allelic imbalance of ultra-rare variants seen in gnomAD against the 504 novel PTVs that were unobserved in gnomAD but present in GTEx (Figure 1B). These novel PTVs showed increased allelic imbalance, indicating that there is not a plateauing of the NMD effect for ultra-rare PTVs.

### NMD efficiency is primarily determined by mutation location, allele frequency, and conservation

The primary means of detecting NMD has been the 50nt rule in which a transcript will be degraded if the PTV occurs upstream of the point 55 to 50 nucleotides prior to the last exon-exon junction (Popp and Maquat 2016; Nagy and Maquat 1998). However, as the 50nt rule alone is not a perfect predictor of NMD efficiency (Supplementary Figure 1), we wanted to investigate if there were more subtle regulatory, tissue-specific, or inter-individual effects that could be detected using the multi-tissue, population design of GTEx. We chose to use the set of predictors previously described in Rivas et. al. 2015 as they had been shown to have predictive power for NMD. Motivated by our previous findings (Figure 1B) and further leveraging the unique capabilities of GTEx, we added gnomAD MAF, tissue, and subject as predictors in the models to test their effects on NMD efficiency.

Initially, we constructed our model to predict allelic imbalance as defined by the binary classification of proportion of reference reads greater than or equal to 0.65 or less than 0.65. Notably, we found that including tissue as a predictive variable did not significantly improve the model (Figure 2A), and including individuals as a predictive variable actually decreased performance (Supplementary Figure 2). Although we did see suggestive evidence for differences in median NMD efficiency between tissues (Supplementary Figure 3), modeling tissue did not increase predictive performance when combined with other information about the PTV. This is similar to what was observed in Rivas et. al., where some samples showed differences in NMD efficiency. Despite our increase in sample size, we were not able to identify a systematic pattern.

**Figure 2.**
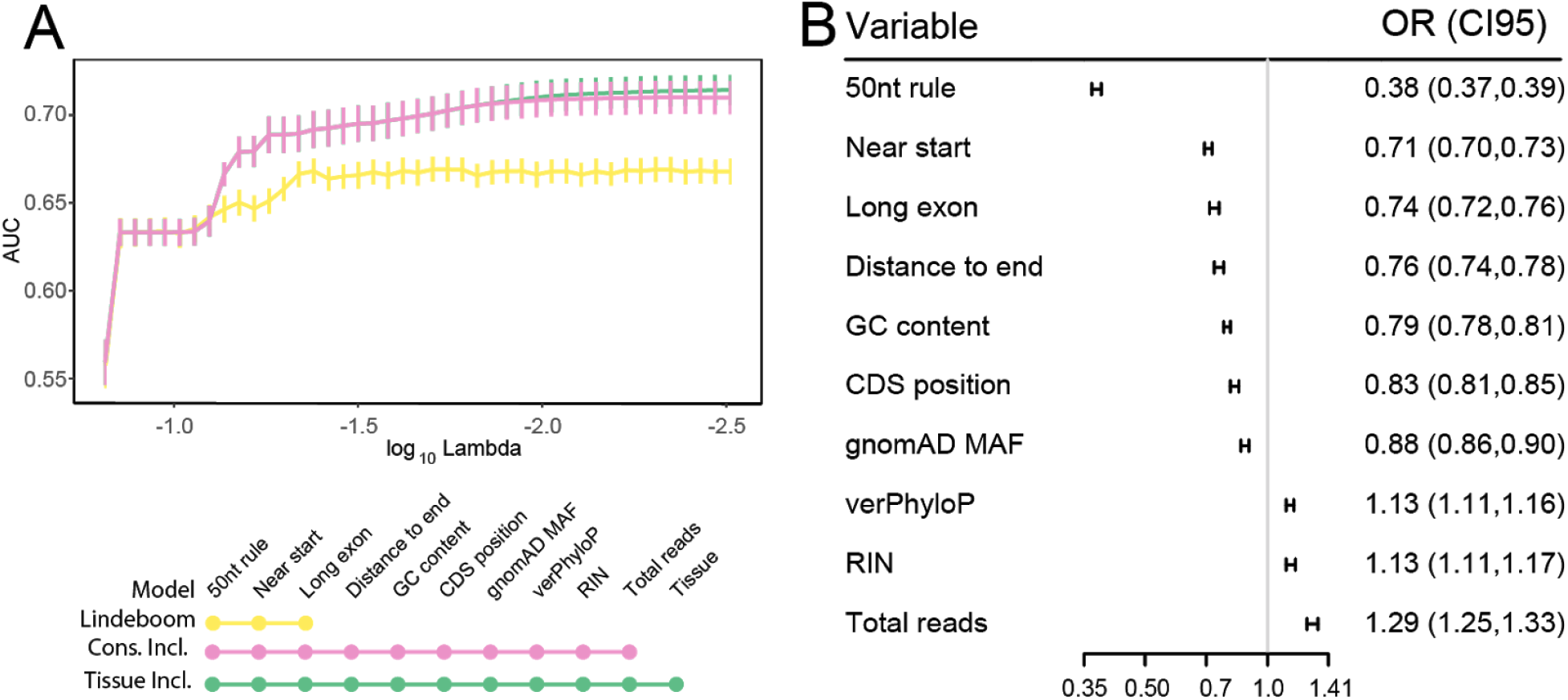
Predictive ability is improved by using variant allele frequency and conservation, but not tissue or subject information. **A**. Plot of model performance over LASSO regularization paths with different feature sets predicting the binary classification of proportion of reference reads ≥ 0.65 or < 0.65. The x-axis shows the log10 value of the regularization parameter lambda, with smaller values corresponding to less penalization. The y-axis shows the area under the curve (AUC) metric of classification performance for each value of lambda. Error bars are obtained from leave-group-out cross validation where the model was trained on all but one chromosome and tested on the left out chromosome. “Lindeboom” model includes 50nt rule, long exon and near start (yellow). “Cons(ervation) Incl(uded)” model also includes the distance to the end (canonical stop), GC content, position in the coding sequence (from start), gnomAD allele frequency, vertebrate phyloP score, RNA integrity number, and total read depth at the site of interest (pink), “Tissue Incl(uded)” adds tissue to the conservation included model (green). **B**. Forest plot of effect sizes and p-values for features that were chosen by the model with the optimal lambda penalty value as measured by AUC using the multi-tissue moderate ASE outcome.

We were further able to leverage the multi-tissue design of the GTEx project to improve performance by predicting the incidence of allelic specific expression (MODASE, equal to 1 - [probability of no ASE]) using a Bayesian stratification approach that reduces noise by including information from multiple observations of a PTV in one individual across tissues (Rivas et al. 2015). Given the integration of multiple tissue information, this approach may reduce our ability to detect tissue specific differences. We proceeded to use MODASE because tissue was not a predictive variable of the proportion of reference reads and the noise reduction led to an improvement in our predictive power (Supplementary Figure 2).

In order to disentangle the often correlated biological predictors, we chose to use the LASSO penalized logistic regression model implemented by the R package *glmnet* to produce a sparse and interpretable model (Friedman, Hastie, and Tibshirani 2010). In addition to the canonical 50nt rule, long exon, and start proximal predictors identified by Lindeboom et. al., we found that the distance to the canonical stop, GC content, position in the coding sequence (the distance from the start), gnomAD allele frequency, vertebrate phyloP score, RNA integrity number, and total read depth at the site of interest were significant predictors of MODASE status (Figure 2B). Unsurprisingly, NMD was easier to detect in samples with higher RNA quality, as denoted by RNA integrity number, and for variants with higher read count.

In order to test the impact of additional factors, we included additional variant and subject level information in our model. Sample level variables that were dropped from the model include: age, sex, cause of death (Hardy scale), and post mortem interval.

### Allelic imbalance of PTVs is consistent across tissues

Based on our observations that tissue was not predictive of allelic imbalance for PTVs, we wanted to evaluate the consistency of allelic imbalance for PTVs across tissues and within an individual. For each individual subject that had the same variant expressed in multiple tissues, we performed a pairwise correlation of the allelic ratio of that variant in those tissues. We were able to investigate most tissue combinations, but we did not have individuals that were sampled for both male-specific tissues (prostate and testis) and female-specific tissues (ovary and vagina) or in two of the lower sampled tissues analyzed (Small Intestine - Terminal Ileum and Brain - Amygdala). We also computed intra-individual, cross-tissue pairwise correlations of a variant’s allelic ratios for missense and synonymous coding variants and non-coding variants in introns, untranslated regions (UTRs) and non-coding exons (Supplementary Figure 4, summarized in Figure 3).

**Figure 3.**
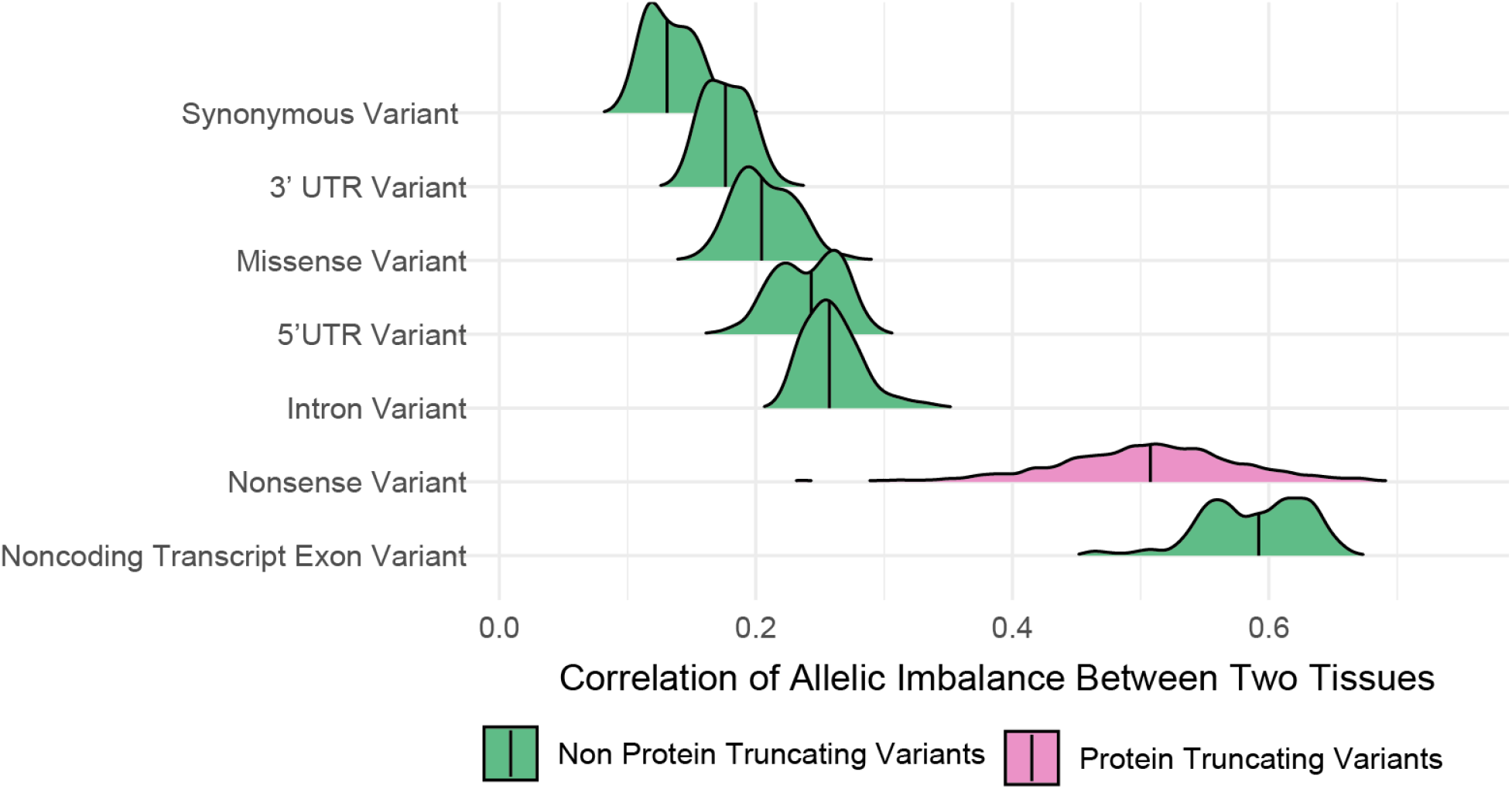
Nonsense variants show more consistent allelic imbalance between pairs of tissues than variants in other coding transcripts. Densities of Pearson correlations of proportion of reference reads for a variant in the same individual in different pairs of tissues. Vertical lines denote median correlation. PTVs are highlighted in pink.

We found a significantly stronger correlation between the proportion of reference reads for all PTVs, with a median Pearson correlation of 0.508, than for any other coding transcript variant, with a median Pearson correlation of 0.131 for synonymous variants and 0.204 for missense variants, or non-coding variants in introns or UTRs (median correlation of 0.176 for 3’ UTRs, 0.243 for 5’ UTRs, and 0.257 for intronic variants). Additionally, PTVs that were predicted to escape NMD showed lower correlation (median 0.449) than those that were predicted to undergo NMD (median 0.552), suggesting the consistency of NMD across tissues.

Intriguingly, noncoding transcripts were the only transcripts that showed higher between-tissue allelic correlations. This higher correlation was not attributable to a systematic difference in read depth (Supplementary Figure 5) or the distribution of the proportion of reference reads between noncoding transcripts and other gene biotypes (Supplementary Figure 6).

### Unpredicted allelic balance is consistent across tissues

Using these models to predict the efficiency of NMD and the RNA sequencing data to verify the effects, we were able to discern which PTVs showed unpredicted allelic balance -- that is, PTVs which are predicted to undergo NMD and are therefore expected to show allelic imbalance but instead show allelic balance. Of the 2,320 rare variants, we found 23.4% (543) variants that showed unpredicted allelic balance at least once, with 7.5% (173) variants showing unpredicted allelic balance in over 90% of observations.

One of the most consistent unpredicted allelic balance variants is rs141826798 in EGFL8 for which we observe the variant in 7 individuals and 44 different tissues for 131 total observations and 98% (128 of the 131) of the observations have a proportion of reference reads below 0.65 (Figure 4A). Inspection of the PTV location shows that it is not near the start, in a long exon, or after 50nt before the last exon junction (Figure 4B) and is not associated with any aberrant splicing events. The variant, rs141826798, was further previously identified as a risk variant for psoriasis in a UK BioBank genome wide association study (Emdin et al. 2018). This further suggests that identification of unpredicted allelic balance can be identified empirically by performing RNA sequencing from a patient of a readily accessible tissue or cell line.

**Figure 4.**
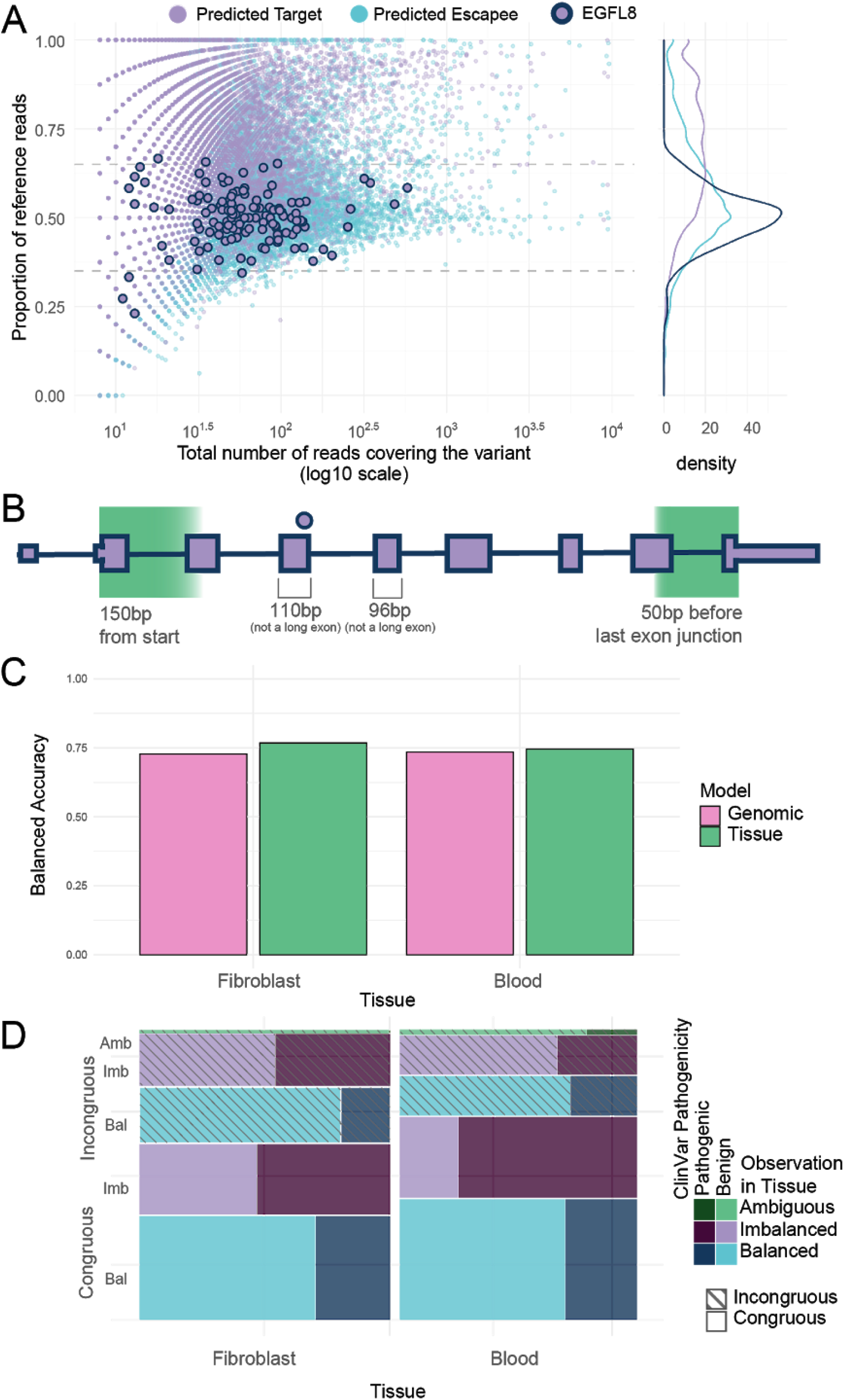
Additional disease relevant information may be gathered by analyzing readily available tissues. **A**. Proportion of reference reads against the total number of reads covering the variant for predicted NMD targets (light purple fill) and variants predicted to escape NMD (light blue fill). The variant rs141826798 in Epidermal Growth Factor Like Domain Multiple 8 (EGFL8), which has been implicated in psoriasis, is highlighted with a navy outline. **B**. The premature termination variant rs141826798 in *EGFL8* occurs 3bp from the end of exon 4. It is not in a long (>407bp) exon, proximal to the start of the gene, in the last exon or 50nt before the last exon junction. **C**. Balanced accuracy of the predictive ability in all other tissues of variants observed in whole blood or fibroblasts using our best predictive model utilizing genomic annotations (pink) or the classification called by the majority of observations across individuals in the indicated tissue (green). **D**. Mosaic plot of the counts of pathogenic (dark colors: dark purple, dark blue, dark green) and benign (light colors: light purple, light blue, light green) variants as determined by ClinVar for variants observed to be all balanced (blues), all imbalanced (purples), or a mixture of both (greens) in each of the indicated tissues. Hashed fills indicate variants for which our predicted model and observed ASE classification differed (incongruous).

In order to determine if utilizing cell lines or readily accessible tissues could provide improved accuracy for determining NMD in all tissues, we looked at the similarity between the allelic imbalance of variants expressed in an easily accessible tissue and cell line (whole blood and fibroblasts) and all other tissues. The category determined by the majority of observations of a variant in fibroblasts and whole blood was a marginally better predictor than our best predictive model using only genomic data. Using the variants observed in fibroblasts to predict the categorical outcome of the variants in other tissues, we found a balanced accuracy of 0.767 as compared to 0.728 for the prediction for the same variants from our genomic model. The balanced accuracy of the predictions derived from blood was 0.745 as compared to 0.734 for the same variants using the predictions from our genomic model (Figure 4C). We found a similar marginal improvement when using Cohen’s Kappa to measure reliability (0.539 versus 0.458 for fibroblast and 0.496 versus 0.465 for whole blood, Supplementary Figure 7).

Because tissue and cell line observations provided additional information for cross-tissue NMD predictions, we wanted to analyze the pathogenicity of the variants for which predicted and observed ASE classification differed. We observed that all imbalanced variants were more likely to be pathogenic (Figure 4D), possibly because GTEx individuals were not selected based on a specific disease phenotype and many imbalanced pathogenic variants are likely recessive. The variants for which our genomic model and the observed classification were congruous (i.e. both imbalanced) showed the largest proportion of pathogenic variants (p<0.0001 Fisher’s exact test for each category).

## Discussion

We analyzed rare protein truncating variants (PTVs) across individuals and tissues using GTEx v8 project data. Using RNA-seq-based measurements of allelic imbalance of PTVs as a measure for NMD efficiency, we observed that, in addition to the position of the variant in the transcript, both allele frequency and conservation were predictive of NMD efficiency. Previous studies have demonstrated increased allelic imbalance of rare versus common PTVs (Kukurba et al. 2014; Rivas et al. 2015; Lappalainen et al. 2013). We observed that these effects did not plateau in the rare portion of the allele frequency spectrum; ultra-rare (MAF < 0.001%) and novel PTVs showed evidence of increased NMD efficiency. Additionally, we observed that GC content impacted NMD efficiency, suggesting an additional role of RNA structure. By combining these factors, we were able to improve our ability to predict whether a PTV would show allelic imbalance beyond the 50nt rule.

Strikingly, NMD efficiency is highly consistent across tissues and individuals, indicating the fundamental importance of this cellular machinery. This is consistent with previous observations of when the NMD machinery fails: individuals with mutations in one of the key NMD factors, UPF3B, show severe intellectual disability and the variant only persists in these families because it is X-linked (Laumonnier et al. 2010; Tejada et al. 2019). As the GTEx individuals were not selected for any phenotypic abnormality, we did not observe any missense or nonsense mutations in the core NMD proteins which may have otherwise provided variation in NMD efficiency between individuals. The relative lack of variation between tissues may be attributed to a finely tuned autoregulatory feedback loop as several of the core NMD proteins are known to be upregulated when NMD is inhibited (Yepiskoposyan et al. 2011).

For classifying novel variants, especially for rare disease diagnostic purposes, it is very promising that NMD is consistent across tissues, age, and sex. The high tissue-sharing of NMD efficiency further indicates that potential gain-of-function effects of NMD escapees, such as those reported by Coban-Akdemir et al, are unlikely to manifest in a single tissue when the target gene is expressed across multiple tissues (Coban-Akdemir et al. 2018). This provides confidence that tissue-agnostic predictive tools such as NMDetectiveB (Lindeboom et al. 2019) provide equal predictive power regardless of the tissue in which a gene is expressed. Further, since tissue is not strongly predictive of NMD efficiency, future studies may benefit from testing in easily biopsied tissues or synthetically testing PTVs in cell lines, with the exception of PTVs in genes with tissue specific splicing. This is especially valuable given the importance of collecting high quality RNA with high coverage at the site of interest. Given datasets like GTEx, it is possible to assess the degradation of many rare variants for appropriate classification without further experiments. To this end, we provide the classification for rare variants identified in this study (Supplementary Table 1) for future research.

## Methods

### Calling nonsense-mediated decay from allele-specific expression

The set of calls generated from BAMs aligned with STAR using the WASP method for allelic mapping bias (van de Geijn et al. 2015) were used. Allele-specific expression (ASE) was called from GTEx v8 data using *ASEAlleleCounter* from GATK (Castel et al. 2015). Nonsense-mediated decay (NMD) from ASE calls was defined as occurring at a protein-truncating variant (PTV) if the ratio of reference reads to the total number of reads was greater than 0.65.

### Variant annotation

Variants were annotated using Variant Effect Predictor (Karczewski et al. 2020) with Ensembl version 88, the same annotation used for other analyses in GTEx v8, except to obtain gnomAD allele frequencies, for which version 97 of the Ensembl annotation was used. The LOFTEE plugin for VEP (Karczewski et al. 2020) was used to obtain the 50nt rule and loss of function prediction. Conservation scores and GC content were obtained from CADD (Kircher et al. 2014; Rentzsch et al. 2019). The canonical isoform was used to select a single annotation for each variant, with variants which were annotated to multiple genes, had multiple predicted consequences, or were annotated as intergenic being removed. NMD was only considered for variants which were exclusively annotated as “stop_gained”. The additional categories “missense_variant”, “synonymous_variant”, “intron_variant”, “3_prime_UTR_variant”, “5_prime_UTR_variant”, and “non_coding_transcript_exon_variant” were used for the comparison of different classes of variants described below.

### Multi-tissue allele-specific expression

The proportion of reference reads does not account for the number of reads supporting it, nor does it exploit the availability of gene expression from multiple tissues in a subject. We used a procedure described in Rivas et. al. 2015 to integrate this information and compute a probability of ASE (Rivas et al. 2015). This normally produces probabilities for no ASE, moderate ASE, or strong ASE. We disabled the estimation of strong ASE in each tissue as this usually exhibited a low probability, that is, a variant was unlikely to be predicted to undergo strong ASE. Using only the moderate ASE measurement provided one probability for the presence of ASE per sample. For every variant, an individual probability was given for each tissue from an individual. NMD was defined as occurring in a given variant if the ASE probability was greater than 0.8 and the proportion of reference reads was greater than 0.5 (to remove variants that exhibited a bias towards the alternate allele).

### Predictive models

NMD efficiency was predicted as either a categorical outcome, using the proportion of reference reads or the multi-tissue ASE probabilities, or a continuous outcome, using L1-penalized regression with the *glmnet* R package (Friedman, Hastie, and Tibshirani 2010) on scaled and centered predictors. For the categorical outcomes, the logistic family was used, with area under the curve (AUC) used as the performance metric, while for the continuous outcomes, the Gaussian family was used on the logit-transformed probabilities, with a correction applied (Smithson and Verkuilen 2006) to adjust probabilities of 0.0 or 1.0, and root mean square error (RMSE) as the performance metric. The penalization parameter lambda was optimized across a range of values chosen by *glmnet* using cross-validation using chromosomes as the folds to guarantee that all variants in the test set were never used in the training set. The best lambda was chosen as the value for which the mean performance metric was best across all 22 folds. P-values and confidence intervals were estimated using the *selectiveInference* package (Tibshirani et al. 2016), using the optimal lambda identified in the cross-validation.

### Correlation of ASE across tissues

We estimated the Pearson’s correlation of ASE across pairs of tissues by matching all instances of a variant being shared between two tissues in the same subject, and correlating the corresponding proportions of reference reads. Separate correlations were performed for each of the following classes of variants: stop gain, missense variant, synonymous variant, intronic variant, non-coding transcript exon variant, 5’ UTR variant, or 3’ UTR variant.

### Assessing NMD variants using ClinVar Pathogenicity information

PTV pathogenicity was assessed using the September 2020 release of the ClinVar Variant Summary table. Data was accessed from https://ftp.ncbi.nlm.nih.gov/pub/clinvar/tab_delimited/variant_summary.txt.gz on September 6, 2020 and filtered for premature termination variants. We then intersected the variants with ClinVar pathogenicity information with the NMD variants recovering 309/2,320 (13%). This accounted for 157 out of 1,189 observations (13.2%) in fibroblasts and 164 out of 1,013 observations (16.2%) in whole blood.

## Acknowledgments

NAT is supported by NIH grants DK107437, HL142015, DK112348, and the Stanford School of Medicine Department of Pathology. DN is supported by 1T32AG047126-01. TE is supported by NLM training grant LM007033. NMF is supported by a National Science Foundation Graduate Research Fellowship, grant no. DGE – 1656518 and a graduate fellowship from the Stanford Center for Computational, Evolutionary and Human Genomics. CS is supported by NIH grant T32LM012409. M.A.R. is in part supported by the NHGRI of the NIH under award R01HG010140 (M.A.R.) and an NIH Center for Multi- and Trans-ethnic Mapping of Mendelian and Complex Diseases grant (5U01 HG009080). SBM is supported by NIH grants R01AG066490, U01HG009431, R01HL142015, R01HG008150, and U01HG009080. This work in-part used supercomputing resources provided by the Stanford Genetics Bioinformatics Service Center, supported by National Institutes of Health S10 Instrumentation Grant S10OD023452. Thanks to the members of Montgomery and Rivas Labs and Sarah Teran for their support and critical feedback in the preparation of this manuscript. The authors thank the GTEx and UKBB participants and their families.

## Supporting information captions

**Supplementary Figure 1.**
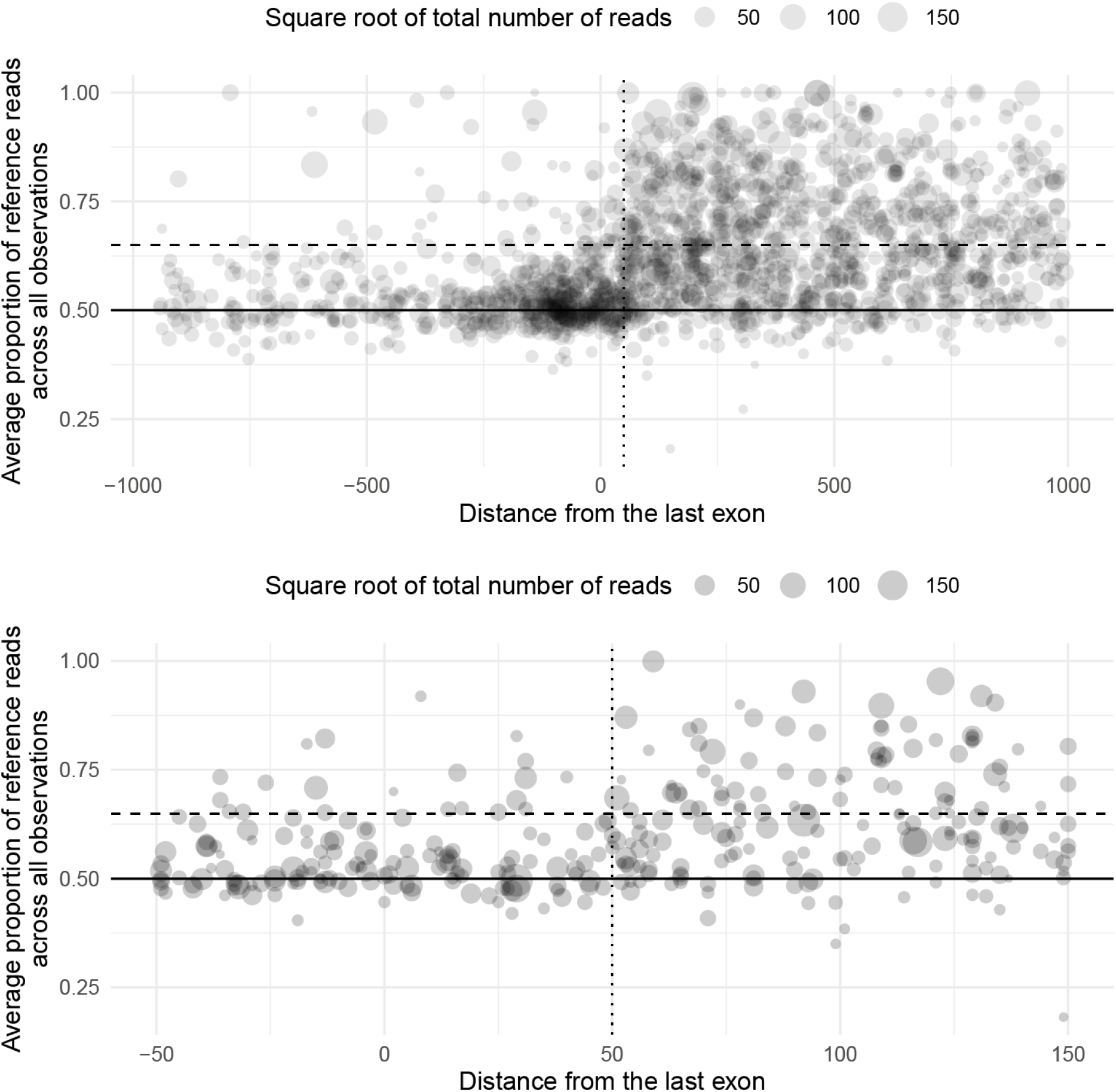
The 50nt rule is an imperfect predictor of whether a variant will be targeted by nonsense mediated decay (NMD). **A**. Scatter plot of the average proportion of reference reads across all observations of a variant, across all individuals and tissues, against the position of that variant relative to the last canonical exon junction. Dot size corresponds to the square root of the total number of reads observed over the variant site, showing only variants from 1000 nucleotides before the last exon junction to 950 nucleotides after. The solid horizontal line shows the expected proportion of reference reads without NMD, 0.5, while the dashed line shows our cutoff for binning a variant as targeted by NMD, 0.65. The vertical dotted line shows 50 nucleotides before the last exon junction. **B**. As in A, but showing only variants from 150 nucleotides before the last exon junction to 50 nucleotides after.

**Supplementary Figure 2.**
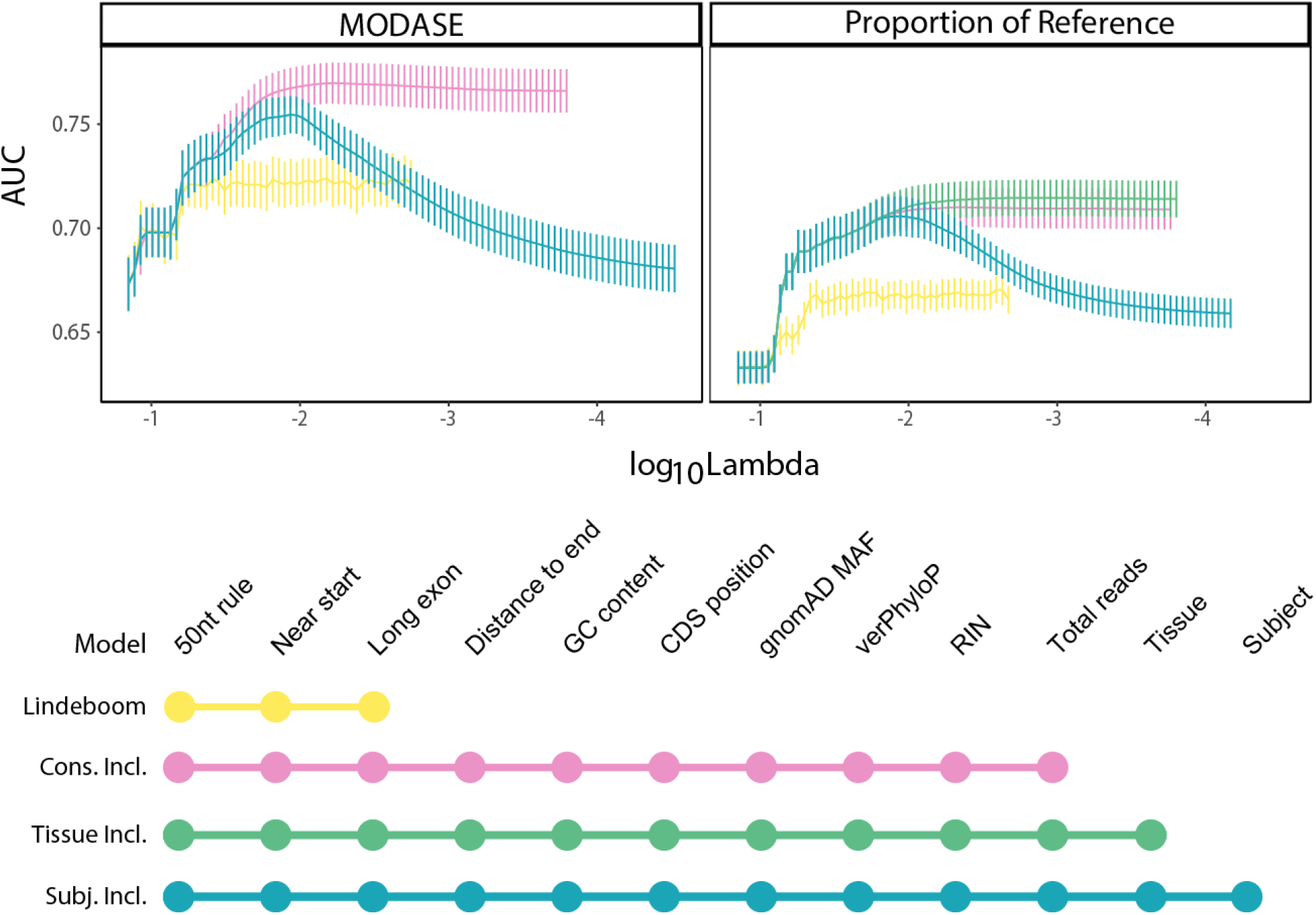
MODASE improves predictive performance over using the measured proportion of reference reads. Plot of model performance over LASSO regularization paths with different feature sets. The x-axis shows the log10 value of the regularization parameter lambda, with smaller values corresponding to less penalization. The y-axis shows the area under the curve (AUC) metric of classification performance for each value of lambda. Error bars are obtained from leave-group-out cross-validation where the model was trained on all but one chromosome and tested on the left out chromosome. “Lindeboom” model includes 50nt rule, long exon and near start (yellow), “Cons(ervation) Incl(uded)” model also includes the distance to the end (canonical stop), GC content, position in the coding sequence (from start), gnomAD allele frequency, vertebrate phyloP score, RNA integrity number, and total read depth at the site of interest (pink), “Tissue Incl(uded)” adds tissue to the conservation included model (green). The left shows the AUC for predicting the MODASE while the right shows the same for the Proportion of Reference reads.

**Supplementary Figure 3.**
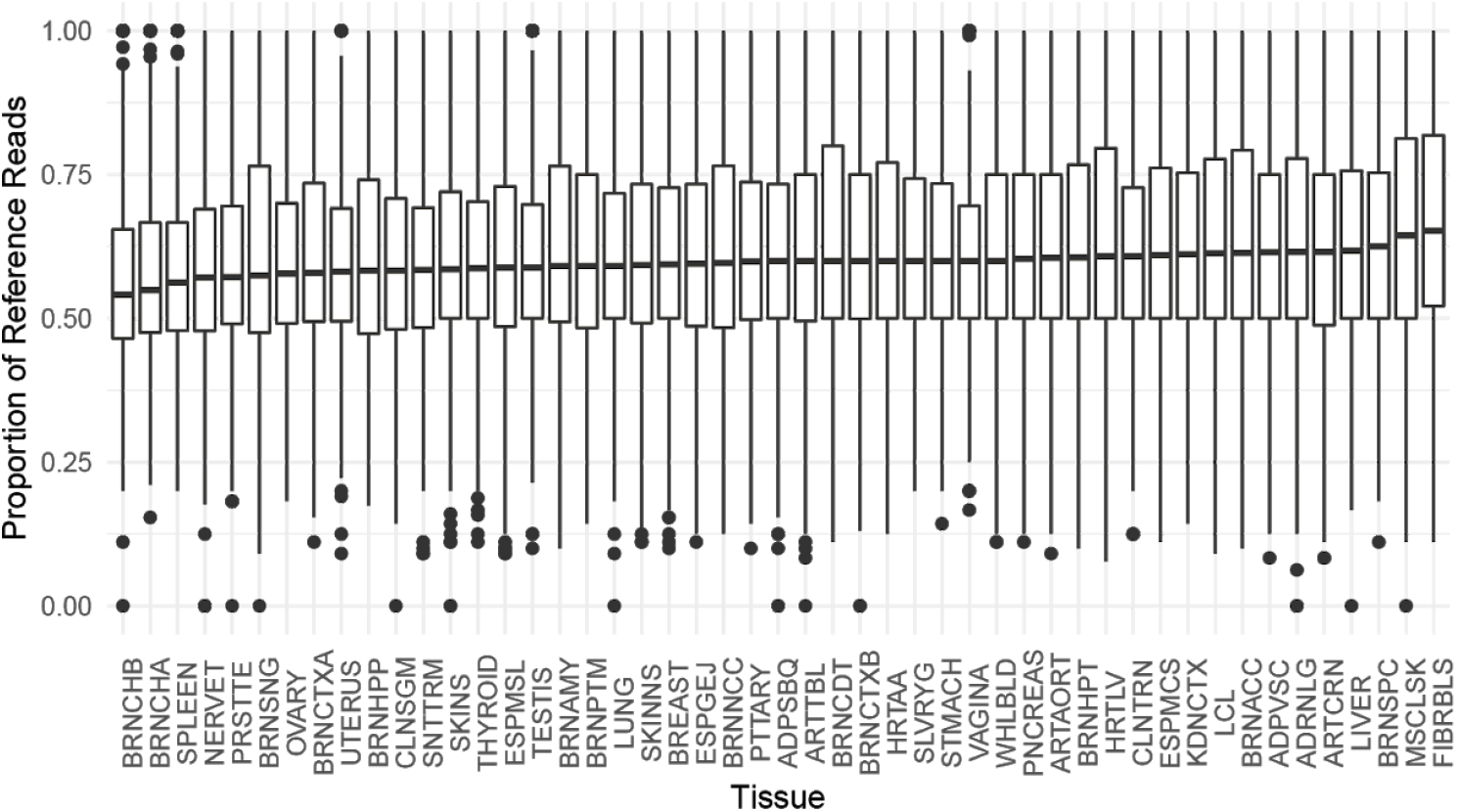
The proportion of reference reads across a protein truncating variant site remains consistent across tissues. Box plot of the proportion of reference reads for each expressed PTV in each tissue. Thick horizontal line shows the median, the box shows the interquartile range, the whiskers show 1.5 times the interquartile range, and the dots show sites beyond the whiskers.

**Supplementary Figure 4.**
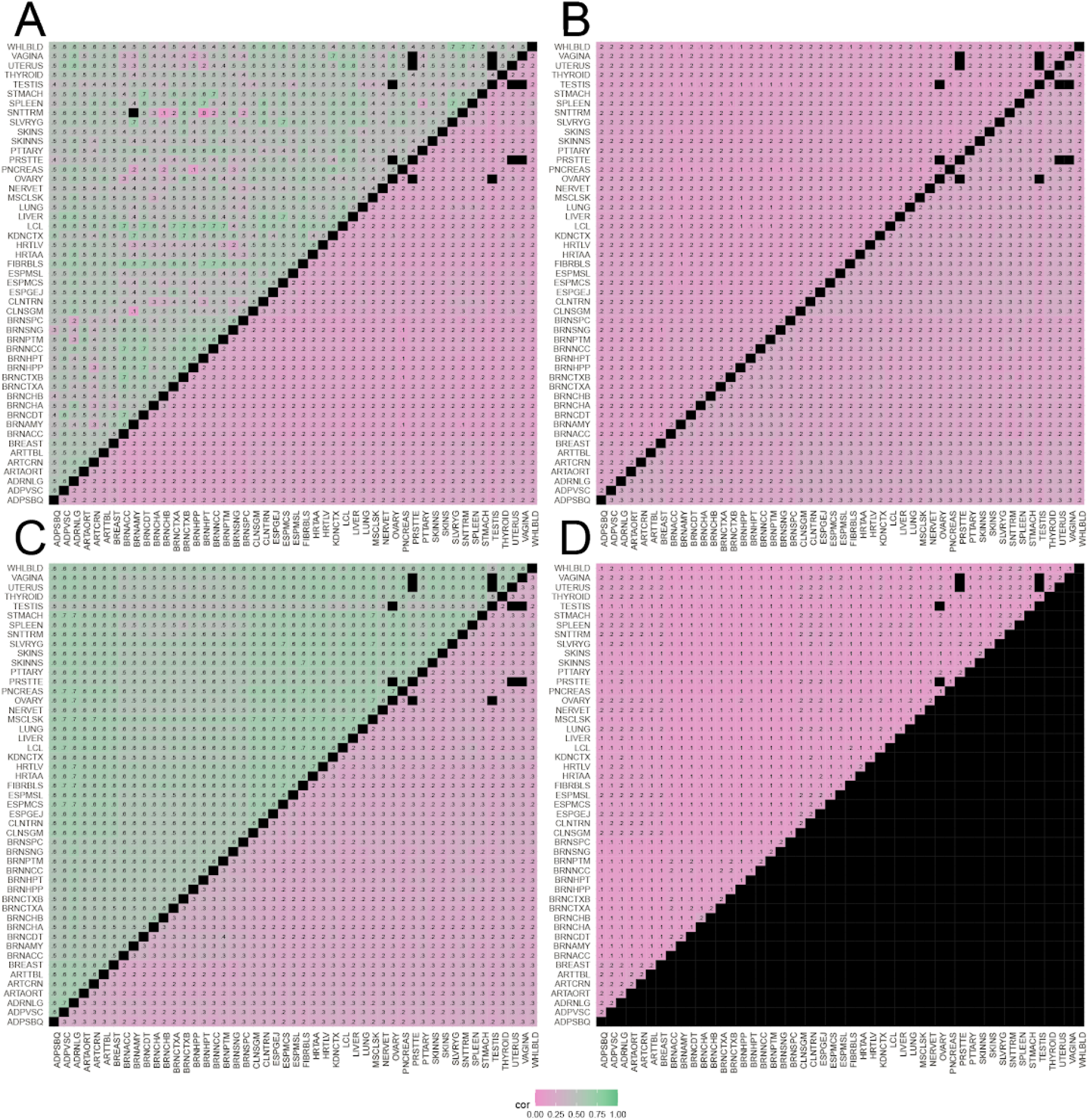
Nonsense variants show more consistent allelic imbalance between pairs of tissues than missense variants. **A**. Pearson correlations of reference allele proportion for PTVs (upper triangle) or missense variants (lower triangle) in the same individual in different pairs of tissues. High correlation: green; low correlation: pink; matrix diagonal or no subject overlap: black. **B**. As in A, but upper half is 3’ UTR variants and lower half is 5’ UTR variants **C**. As in A, but upper half is non-coding transcript exon variants and lower half is intronic variants **D**. As in A, but upper half is synonymous variants.

**Supplementary Figure 5.**
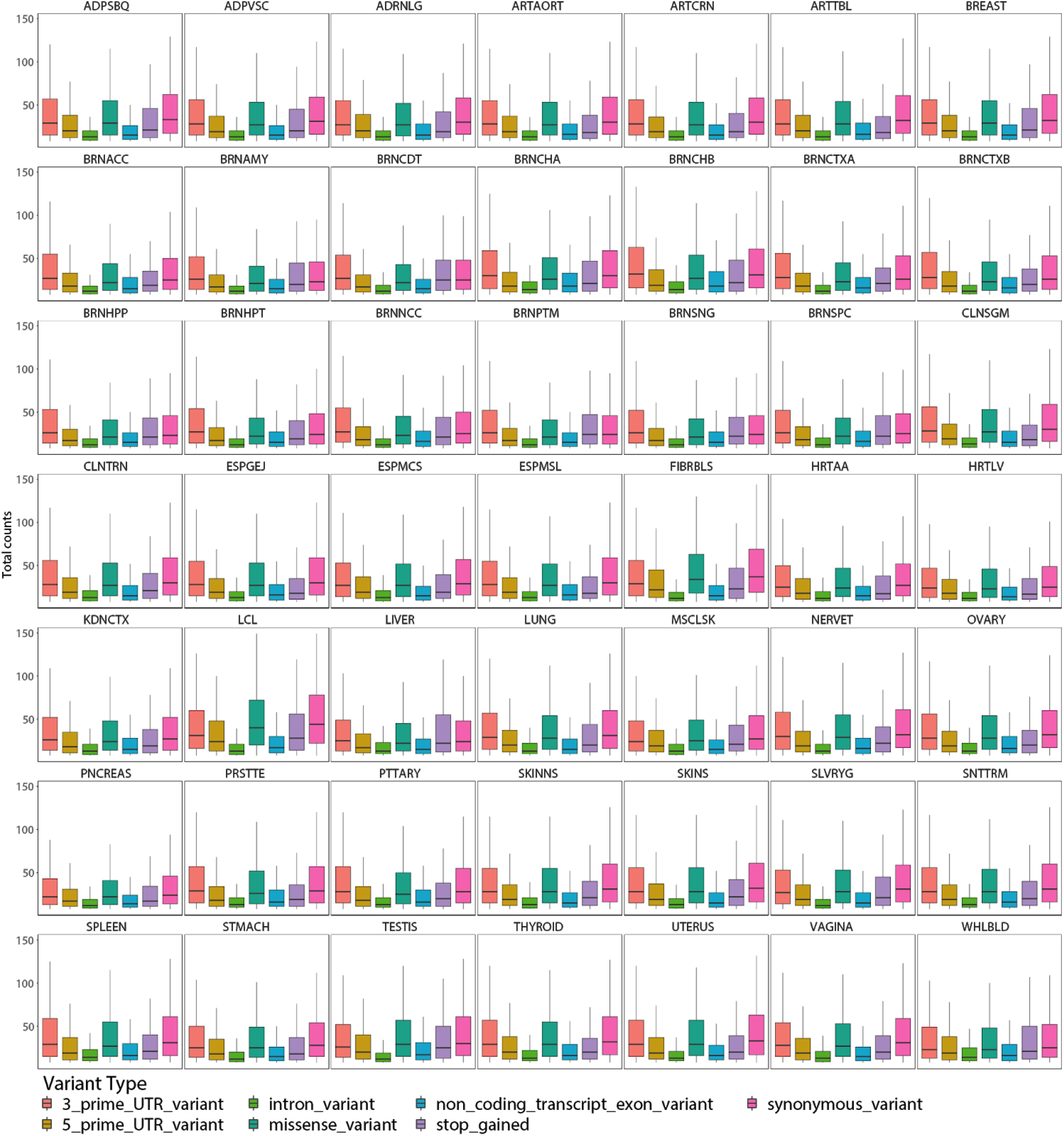
ASE variants in non-coding transcript exons have a comparable number of counts to variants in other genomic regions. For each tissue, a box plot summarizes the distribution of total read counts of ASE variants in each genomic region. A small number of variants had an extremely high number of reads. To aid in visualization, variants with more than 200 reads are not shown.

**Supplementary Figure 6.**
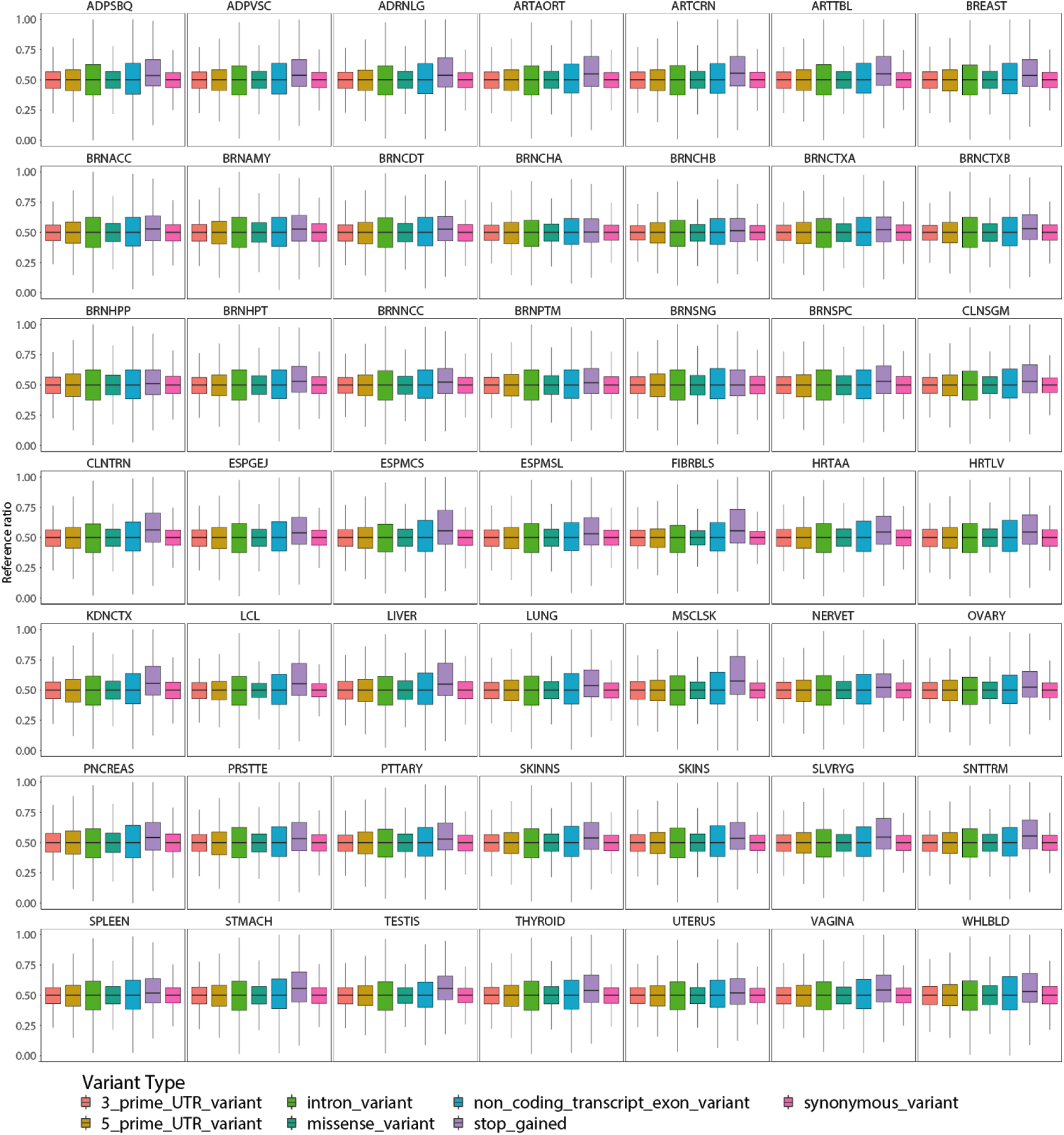
ASE variants in non-coding transcript exons have a comparable distribution of reference ratios to other genomic regions. For each tissue, a box plot summarizes the distribution of reference ratios in ASE variants in each genomic region.

**Supplementary Figure 7.**
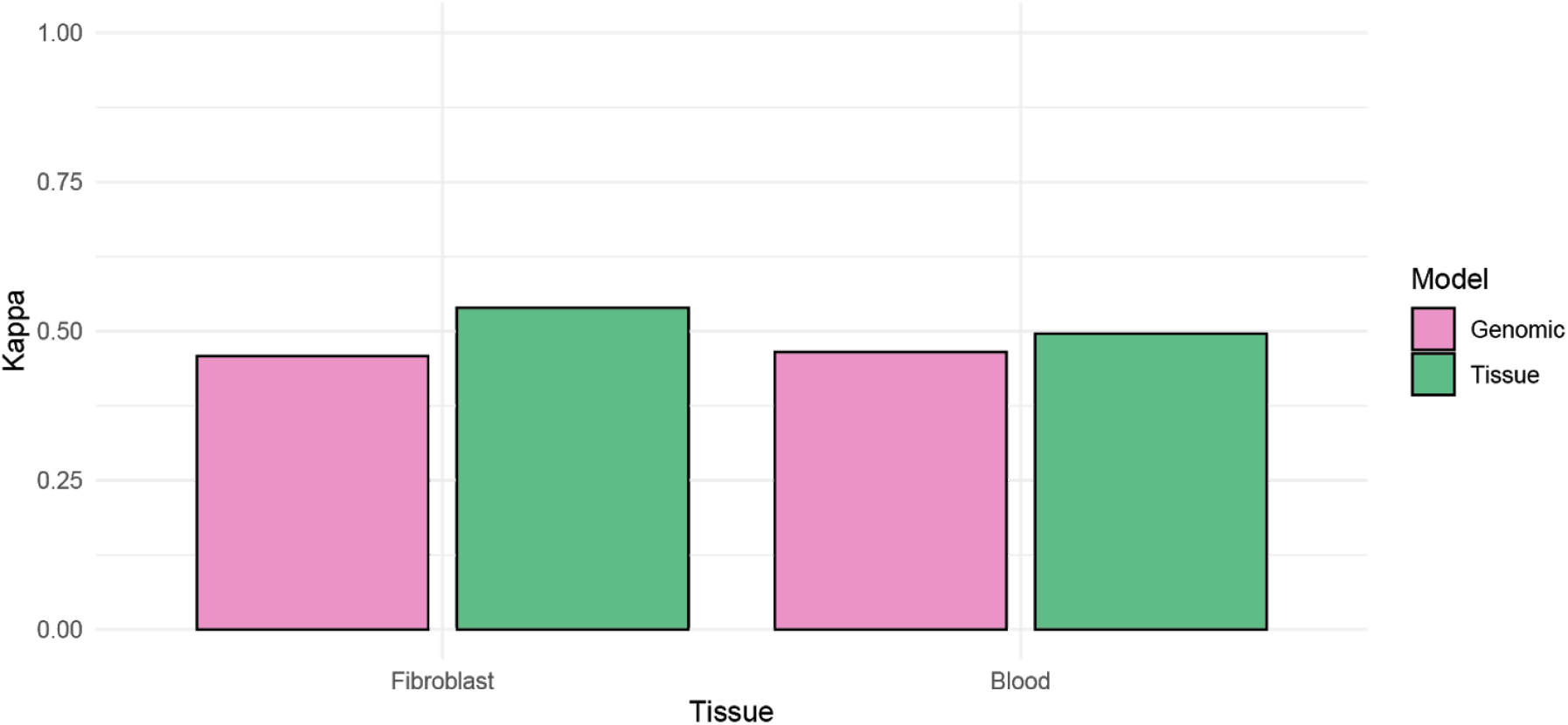
Additional disease relevant information may be gathered by analyzing readily available tissues. Cohen’s Kappa of the predictive ability in all other tissues of variants observed in whole blood or fibroblasts using our best predictive model using genomic annotations (pink) or the classification called by the majority of observations across individuals in the indicated tissue (green).

